# Demography and natural selection have shaped genome-wide variation in the widely distributed conifer Norway Spruce (*Picea abies*)

**DOI:** 10.1101/805903

**Authors:** Xi Wang, Carolina Bernhardsson, Pär K. Ingvarsson

## Abstract

Under the neutral theory, species with larger effective population sizes are expected to harbour higher genetic diversity. However, across a wide variety of organisms, the range of genetic diversity is orders of magnitude more narrow than the range of effective population size. This observation has become known as Lewontin’s paradox and although aspects of this phenomenon have been extensively studied, the underlying causes for the paradox remain unclear. Norway spruce (*Picea abies*) is a widely distributed conifer species across the northern hemisphere and it consequently plays a major role in European forestry. Here, we use whole-genome re-sequencing data from 35 individuals to perform population genomic analyses in *P. abies* in an effort to understand what drives genome-wide patterns of variation in this species. Despite having a very wide geographic distribution and an enormous current population size, our analyses find that genetic diversity of *P.abies* is low across a number of populations (π=0.005-0.006). To assess the reasons for the low levels of genetic diversity, we infer the demographic history of the species and find that it is characterised by several re-occurring bottlenecks with concomitant decreases in effective population size can, at least partly, provide an explanation for low polymorphism we observe in *P. abies*. Further analyses suggest that recurrent natural selection, both purifying and positive selection, can also contribute to the loss of genetic diversity in Norway spruce by reducing genetic diversity at linked sites. Finally, the overall low mutation rates seen in conifers can also help explain the low genetic diversity maintained in Norway spruce.

## Introduction

Explaining the distribution of genetic diversity within and between species is one of the major goals of evolutionary biology (Begun et al. 2007; Wang et al. 2016). This question has several important applications (Ellegren and Galtier 2016) including the conservation of endangered species (Forcada and Hoffman 2014), and development of breeding strategies in crops (Khodadadi et al. 2011) and farm animals (Edea et al. 2015). Ultimately, genetic diversity reflects the balance between the appearance and disappearance of unique variants. Spontaneous mutation give rise to new variants and under the neutral theory, the ultimate loss or fixation of these new variants is governed by genetic drift (Kimura 1983). Therefore, species with larger effective populations are expected to harbour higher genetic diversity due to a greater influx of mutations and smaller effects of genetic drift. However, observations suggest that the range of genetic diversity among different organisms is orders of magnitude more narrow than the corresponding range of effective population sizes (Leffler et al 2012). This long-standing issue in evolutionary biology is known as ‘Lewontin’s paradox’ (Lewontin 1974; Leffler et al. 2012).

One potential resolution to Lewontin’s paradox is the influence from demographic fluctuations (Ellegren and Galtier 2016) as a consequence of various historical events, such as glacial periods (Leite et al. 2016), geological events (Yan et al. 2013) or human intervention (Qiu et al. 2015). A sharp and rapid decrease in the size of a population (bottleneck), for example, can result in a substantial loss of genetic diversity due to enhanced genetic drift and reduced efficacy of natural selection. Another possible explanation for Lewontin’s paradox is natural selection which reduce levels of neutral polymorphism at sites linked to beneficial mutations under positive selection (“hitchhiking”) (Smith and Haigh 1974) and/or sites under purifying selection acting on deleterious mutations (“background selection”) (Charlesworth et al. 1995; Corbett-Detig et al. 2015). If natural selection is pervasive across the genome of an organism, such linked selection can result in severe reductions of the genome-wide average nucleotide diversity. A hallmark signature of such linked selection is a positive correlation between recombination rate and neutral diversity, as the influence of linkage is expected to be stronger, and will hence remove more neutral polymorphisms, in regions of low compared to high recombination. Such a pattern is unlikely to arise from demographic processes alone (Cutter and Payseur 2013; Corbett-Detig et al. 2015; Ellegren and Galtier 2016;Wang et al. 2016). Reductions in nucleotide diversity associated with selection may also be apparent in regions where the density of selected mutations is high, because such regions are expected to have undergone stronger effects of linked selection (Flowers et al. 2011; Cutter and Payseur 2013; Ellegren and Galtier 2016). Assuming that genes represent the most likely targets of natural selection, linked selection should be stronger in gene dense regions, resulting in an expectation of a negative correlation between gene density and neutral diversity. However, spurious results can be generated when two explanatory variables co-vary but are analysed separately and a positive or negative covariation between recombination rate and gene density are thus expected to either obscure or strengthen the signatures of linked selection across the genome (Flowers et al. 2012; Ellegren and Galtier 2016). Finally, the “drift-barrier” hypothesis states that species with larger populations tend to have lower mutation rates (Lynch et al. 2016). As most spontaneous mutations are likely deleterious, selection should favour lower mutation rates (Xu et al. 2019). Natural selection should also have a stronger effect in larger populations because genetic drift in smaller population overrides the effect of natural selection (Xu et al. 2019) and this thus provides yet another explanation for Lewontin’s paradox.

With the advent of next-generation sequencing (NGS) technologies, unprecedented amounts of genomic data from non-model organism with large populations have made it possible to study the factors contributing to Lewontin’s paradox. Conifers are the most widely distributed group of gymnosperms and are estimated to cover approximately 39% of the world’s forests (De La Torre et al. 2014). Conifers are characterised by an outbreeding mating system with wind-dispersed seeds and pollen, large effective population sizes and extremely large genomes (20-30 Gbp) which contain a large fraction of repetitive DNA, mostly in the form of transposable elements (Nystedt et al. 2013). Estimates of nucleotide diversity in conifers reported so far are surprisingly low, given their life-history traits and widespread distribution (π=0.0024-0.0082 in four conifers from Alpine European forests, *Abies alba*, *Larix decidua*, *Pinus cembra*, *Pinus mugo*, Mosca et al. 2012; π=0.0045 in *Cryptomeria japonica*, Uchiyama et al. 2012; π=0.00132 in *Pinus elliottii* and π=0.00136 in *Pinus taeda*, Acosta et al. 2019). Conifers are thus intriguing organisms for understanding the factors that contribute to low nucleotide diversity and how this can help explain Lewontin’s paradox.

Norway spruce (*P. abies*) is one of the most important ecological and economical conifers and has wide natural distribution that range from the west coast of Norway to the Ural mountains and across the Alps, Carpathians and the Balkans in central Europe (Farjon 1990; Bernhardsson et al. 2019). Earlier studies of nucleotide diversity in Norway spruce have thus far either been limited to coding regions (e.g Heuertz et al 2006, Chen et al. 2012) or to data generated by exome capture sequencing which only covers between 1% to 2% of a typical eukaryote genome (Warr et al. 2015; Baison et al 2019; Chen et al. 2019). However, the recent publication of a reference genome for Norway spruce (Nystedt et al 2013) has opened up possibilities for whole genome re-sequencing in this species and thus makes it possible to assess genome-wide levels of nucleotide diversity. Here we use data from whole genome re-sequencing in a set of tree samples spanning the distribution range of *P. abies* and study how patterns of nucleotide diversity vary across the genome in Norway spruce.

## Materials and Methods

### Sample collection and DNA sequencing

We sampled 35 individuals of Norway spruce (*P. abies*) spanning the natural distribution of the species, extending from Russia and Finland in the east to Sweden and Norway in the west and Belarus, Poland and Romania in the south (Supplementary Table S1; Supplementary Figure S1). Samples were collected from newly emerged needles or dormant buds and stored at −80 °C until DNA extraction. For all samples, genomic DNA was extracted using a Qiagen plant mini kit following manufacturer’s instructions. All sequencing was performed at the National Genomics Initiative platform (NGI) at SciLifeLab facilities in Stockholm, Sweden using paired-end libraries with an insert size of 500bp. Sequencing was performed using a number of different Illumina HiSeq platforms (HiSeq 2000 and HiSeq X). The original location, platform used, estimated coverage from raw sequencing reads and of BAM files after mapping is given for all individuals in Supplementary Table S1.

### Data subsets, SNP calling and hard filtering

Raw sequence reads were mapped to the complete *P. abies* reference genome v.1.0 (Nystedt et al. 2013) using BWA-MEM with default settings in bwa v0.7.15 (http://maq.sourceforge.net/bwa-man.shtml; Li 2013). The Norway spruce genome has an extremely large genome size (∼20Gb) and very large repetitive fraction (∼70%), and analysing genome-wide sequencing data from such massive genomes using a highly fragmented genome assembly (∼10M unique scaffolds) is difficult using existing software. In particular, the limiting variable is the number of genomic scaffolds which is far greater than what existing software can handle. In order to efficiently analyse the data we therefore performed several steps to reduce the computational complexity of the SNP calling process by subsetting data sets into a number of smaller data sets. The smaller subsets enabled us to curate the data in parallel and were crucial for enabling software tools, such as GATK, to function properly.

Following read mapping against the entire reference genome, we reduced the reference genome by only keeping genomic scaffolds greater than 1kb in length using bioawk (https://github.com/lh3/bioawk). All BAM files were then subsetted using the reduced reference genome with the ‘view’ module in samtools v1.5 (http://samtools.sourceforge.net; Li et al. 2009). For each individual, reduced BAM files resulting from different sequencing flow cells were merged into a single BAM file using the ‘merge’ module in samtools and was then split using samtools ‘view’ into 20 independent subsets, where each subset contain roughly 100,000 scaffolds. The reduced reference genome was simultaneously subdivided into the corresponding 20 genomic subsets by keeping the corresponding scaffolds using bedtools v2.26.0 (Quinlan and Hall 2010). The 20 BAM subsets for each individual and 20 reference subsets were finally indexed to make them available for subsequent data processing.

Before SNP calling, PCR duplicates were marked in all data subsets using MarkDuplicates in Picard v2.0.1 (http://broadinstitute.github.io/picard/) to eliminate artefacts introduced due to DNA amplification by polymerase chain reaction (PCR), which could potentially lead to excessively high read depth in some regions. If not addressed, this could bias the number of variants called and may substantially influence the accuracy of the variant detection (Mielczarek and Szyda 2016). Artefacts in alignment, usually occurring in regions with insertions and/or deletions (indels) during the mapping step, can result in many mismatching bases relative to the reference in regions of misalignment. Local realignment was thus performed to minimise such mismatching bases by first detecting suspicious intervals using RealignerTargetCreator, followed by realignment of those intervals using IndelRealigner, both implemented in GATK v3.7 (DePristo et al. 2011). Finally we performed SNP calling using GATK HaplotypeCaller to generate intermediate genomic VCFs (gVCFs) on a per-subset and per-sample basis (20 gVCFs produced for each individual). These gVCF files were then used for joint calling across all 35 samples using the GenotypeGVCFs module in GATK. The SNP calling pipeline produced 20 VCF files where each file contains variants from all 35 individuals for the corresponding subsets.

In order to remove potential false positives in raw variant calls, we performed hard filtering for each of the 20 genomic subset VCF files using the following steps: (1) We only included biallelic SNPs positioned > 5 bp away from an indel and where the SNP quality parameters fulfilled GATK recommendations for hard filtering (https://gatkforums.broadinstitute.org/gatk/discussion/2806/howto-apply-hard-filters-to-a-call-set). (2) In order to reduce the impact from reads mapping to repetitive regions that may be collapsed in the reference genome, we re-coded genotype calls with a depth outside the range 6-30 and a GQ < 15 to missing data and filtered each SNP for being variable with an overall average depth in the range of 8-20 and a ‘maximum missing’ value of 0.8 (max 20% missing data). (3) We removed all SNPs that displayed a *p*-value for excess of heterozygosity less than 0.05, as SNPs called in collapsed regions in the assembly, likely containing non-unique regions in the genome, should show excess heterozygosities as they are based on reads that are derived from different genomic regions. For more detailed information on the genotype calling pipeline and the hard filtering criteria please refer to Bernhardsson et al. (2019). All SNPs that passed the different hard filtering criteria were used in downstream analyses.

### Analysis of relatedness and population structure

We used the relatedness2 option in vcftools (Danecek et al. 2011) to estimate genetic relatedness between all individuals based on the SNP data. This option implements an algorithm for relatedness inference for any pair of individuals that also allows for the presence of unknown population substructure (Manichaikul et al. 2010). During these analyses, two individuals, sampled from same population in northern Sweden, were identified as being half-sibs, and we therefore randomly removed one individual from the pair (Pab034) (Supplementary FigureS2) from subsequent analyses.

To investigate population structure, we performed principal component analysis (PCA) on all SNPs using the module ‘pca’ in plink v1.90. This module extracts the top 20 principal components of the variance-standardized relationship matrix (http://www.cog-genomics.org/plink/1.9/; Chang et al. 2015). A dendrogram was built using Ward’s hierarchical clustering method (Ward 1963; Murtagh and Legendre 2011) as implemented in the ‘hclust’ function in R v3.3.3 (R Core Team 2017) with the pairwise relatedness matrix calculated from method described in Yang et al. (2010) as input. *Picea obovata* (individual Pab001), alternately described as a sub-species (Krutovskii and Bergmann 1995) or sister species (Tsuda et al 2016) of *P. abies*, was used as outgroup in the analyses.

### Estimation of nucleotide diversity

We used ANGSD v0.921 (Korneliussen et al. 2014) to estimate nucleotide diversity separately for all Norway spruce populations. We first used module ‘dosaf 1’ to calculate the site allele frequency likelihood based on normalized phred-scaled likelihoods of the possible genotypes (PL tag in VCF file). The global folded site frequency spectrum (SFS) was than calculated using ‘realSFS’ based on the expectation maximization (EM) algorithm (Nielsen et al. 2012). We then ran the function ‘doThetas 1’ to calculate thetas for each site from the posterior probability of allele frequency (global folded SFS) based on a maximum likelihood approach (Kim et al. 2011). From the per site thetas, pairwise nucleotide diversity (π) (Tajima 1989) per scaffold was final calculated by ‘ThetaStat’.

We repeated the analyses to estimate pairwise nucleotide diversity at different categories of functional sites: 4-fold synonymous sites, 0-fold non-synonymous sites, introns and intergenic sites. Bed files for these different genomic regions were generated from the genome annotation for *P. abies* v1.0 (available from ftp://plantgenie.org/Data/ConGenIE/Picea_abies/v1.0/GFF3/) using a custom-made python script (https://github.com/parkingvarsson/Degeneracy/). Separate VCF files were then generated for the different site categories from the original VCF files based on the genomic bed files using vcftools and were used as input for ANGSD. When computing the global SFS for all SNPs and for intergenic variants in the Sweden-Norway population, we randomly down-sampled number of SNPs to 50% to reduce the computational cost and enable ANGSD to perform analyses.

### Demographic history inference

The demographic history was also inferred based on all SNPs using a coalescent simulation-based method implemented in fastsimcoal2 v2.6.0.3 (Excoffier et al. 2013). We estimated the folded two-dimensional joint site frequency spectrum (2D SFS) to minimize potential bias arising when determining ancestral allelic states using a custom made perl script (https://github.com/wk8910/bio_tools/tree/master/01.dadi_fsc). Eight plausible demographic models were then tested. All models included three current-day populations that were derived from a common ancestral population that experienced an ancient population bottleneck. Following the bottleneck, the Central Europe was assumed to diverge from the ancestral population, followed by the divergence between of the Finnish and Swedish/Norwegian populations (Supplementary Figure S3). The models differed depending on population sizes following divergence and whether individual populations went through further bottlenecks. For each model, we ran 50 independent runs to calculate global maximization likelihood, with 100,000 coalescent simulations per likelihood estimation (-n10000) and 20 conditional maximization algorithm cycles (-L20). The best model was chosen using Akaike’s weight of evidence as described in Excoffier et al. (2013). To obtain the 95% confidence interval of the best model, we generated 100 parametric bootstraps based on the maximum likelihood parameters estimated in best model and ran 50 independent runs for each bootstrap using the same settings as for the analyses of the original dataset. A mutation rate of 8.0×10^-10^ per site per year and a generation time of 25 years was used for *P.abies* (De La Torre et al. 2017; Chen et al. 2019) to convert model estimates from coalescence units to absolute values (i.e. years).

### Correlation between genomic features: nucleotide diversity, divergence, recombination rate, gene density, GC density and repeat density

In order to understand the factors determining variation in genetic diversity across the *P. abies* genome, we estimated correlations between nucleotide diversity at different functional sites (4-fold, 0-fold, introns, intergenic) of each population with different genomic features. The population-scaled recombination rate (ρ=4*N_e_r*) was estimated per scaffold for each population using a Bayesian reversible-jump Markov Chain Monte Carlo scheme (rjMCMC) under the crossing-over model as implemented in LDhat v2.2 (McVean et al. 2004). We performed 1,000,000 MCMC iterations with sampling every 2000 iterations and set up a block penalty parameter of 5 using a data set consisting of only scaffolds longer than 5kb because short scaffolds generally did not produce stable estimates. The first 100,000 iterations of the rjMCMC scheme were excluded as a burn-in. We measured gene density per scaffold as the ratio of sites falling within a gene model on the scaffold to the overall length of the scaffold. The same method was used to estimate repeat density using information on repeat content per scaffold (ftp://plantgenie.org/Data/ConGenIE/Picea_abies/v1.0/GFF3/Repeats/). GC density was calculated at the scaffold level as the fraction of bases where the reference sequence (*P. abies* genome v1.0) was a G or a C using bedtools. Divergence was calculated between each population of Norway spruce with outgroup at 4-fold, 0-fold, intronic and intergenic sites by measuring the number of fixed differences per scaffold. Pairwise correlations between the variables of interest were calculated using Spearman’s rank correlations and by linear regression in Rstudio.

## Results and Discussion

Whole genome re-sequencing data were generated for 35 individuals sampled to span the natural distribution of *P. abies* using Illumina HiSeq 2000/X with a mean sequence coverage of 18.1x per individual (Supplementary Table S1). Raw reads were mapped to the whole genome assembly v1.0 of Norway spruce, which contain ∼10 million scaffolds covering 12.6 Gb out of the estimated genome size of ∼20 Gb (Nystedt et al. 2013). BAM files were reduced to include only scaffolds longer than 1kb (∼2 million scaffolds covering 9.5 Gb of the genome). For each individual, all reduced BAM files were merged into a single BAM file and then were subdivided into 20 genomic subsets with ∼100,000 scaffolds in each. These genomic subsets covered lengths ranging from 159.1 Mb (subset 12) to 2654.8 Mb (subset 1) with an average scaffold length of 4.8 kb (Supplementary Table S2). After read mapping we successively performed PCR duplication removal, local realignment and finally variant calling resulting in 20 raw VCF files containing a total of 749.6 million variants of which 709.5 million were single nucleotide polymorphisms (SNPs) (Supplementary Table S2). A series of stringent filtering criteria were implemented on the raw variant calls to identify high-quality SNPs (see “Materials and Methods” section for details). Following filtration, 293.9 million SNPs remained for all downstream analyses. These SNPs were distributed over 63.2% of the 1,970,460 scaffolds in the reduced assembly reference containing scaffolds longer than 1kb (Supplementary Table S2).

### Population structure

In order to account for effects arising from population structure, we performed a principal component analysis on all SNPs from 34 individuals (Figure1B; Figure1C). Individual Pab001, which is an individual from the sister species to *P. abies*, *P. obovata*, was completely separated from samples of *P. abies* (Figure1B), although previous research have suggested that the two species are genetically quite similar (Krutovskii and Bergmann 1995) and have historically hybridised frequently (Tsuda et al. 2016). We have used the *P. obovata* individual as an outgroup in all downstream analyses in this study where the analyses requires polarisation of variants into ancestral and derived states. After removing the *P. obovata* individual the PCA result clearly reflects geographic structure within *P. abies*, with the top two principle components (PC1-PC2) explaining 1.70% and 1.36% of the total variation in the SNP data, respectively (Figure1C). The individuals of *P. abies* clustered into four groups, consistent with the geographic origin of the samples (Figure 1C; Supplementary Figure S1; Supplementary TableS1): Samples from Belarus, Poland, and Romania grouped into one cluster, most Finish samples clustered into one group, most individuals from Norway and northern Sweden clustered into one group. Two individuals, one from southern Finland (Pab015) and one from southern Sweden (Pab002), could not be grouped within any of the populations as they fell between the three main groups in the PCA plot, indicating population admixture, recent hybridization or that they represent materials that have recently been introduced from elsewhere in Europe. The population structure inferred using the PCA was further supported by a dendrogram constructed using Ward’s hierarchical clustering methods on the pairwise relatedness matrix. Again, individual Pab001 (*P. obovata*) fell outside all *P.abies* individuals and the dendrogram showed that the remaining *P. abies* samples clustered into four groups corresponding to the same groupings inferred in the PCA (Figure1A). In the dendrogram, the Sweden-Norway group clustered together with northern Finish group, while the Central Europe group were closely associated with the two individuals derived from southern Sweden and Finland (Figure1A). Both the results of the PCA and the dendrogram thus suggest a potential population history within Norway spruce where individuals from Central Europe first split from common ancestor, followed by the divergence between Sweden-Norway and Finland.

**Figure 1.**
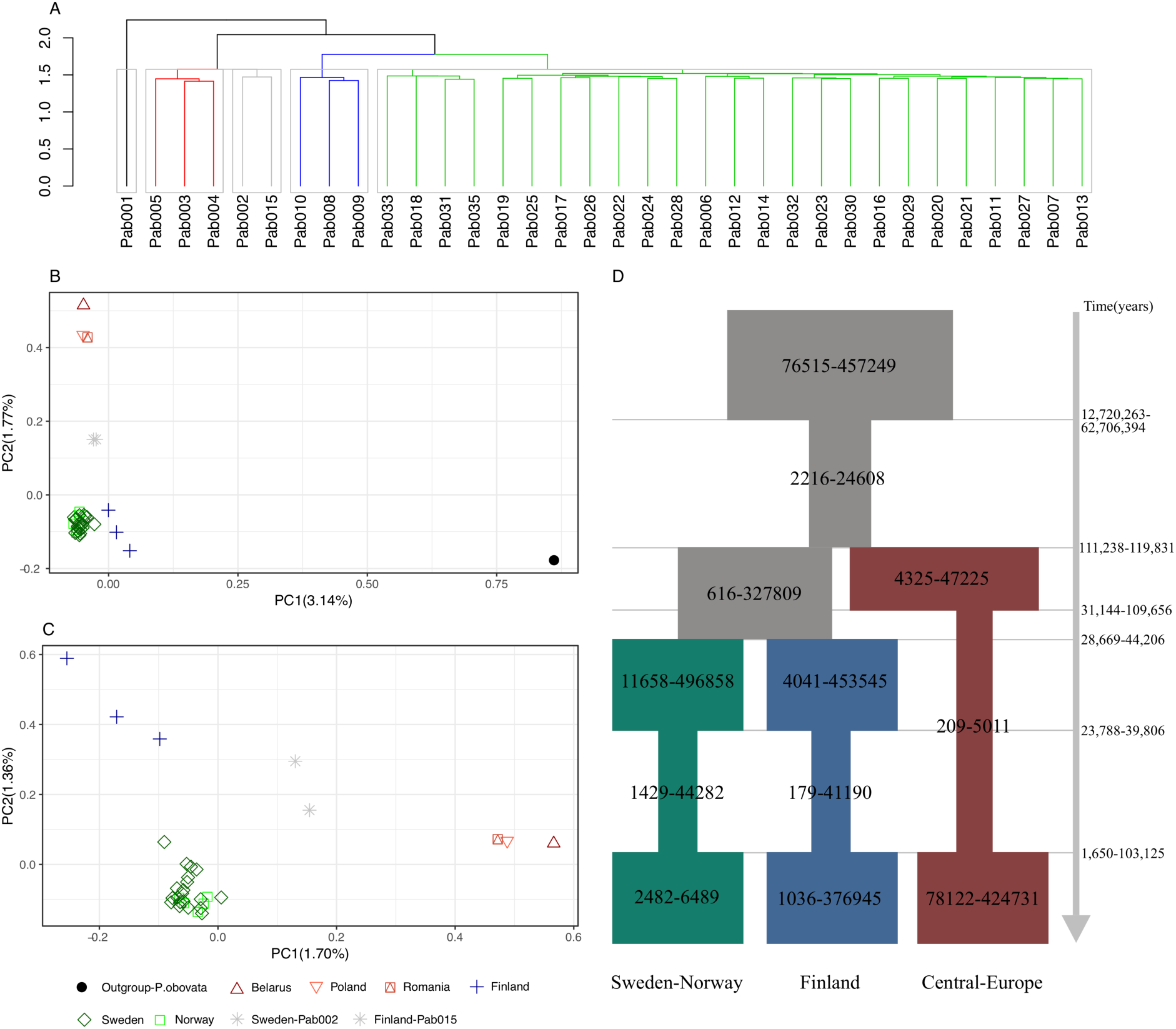
Population structure and demographic history of *P. abies*. (A) The dendrogram was built based on Ward’s hierarchical clustering, using a pairwise relatedness matrix for all individuals. The coloured branches and grey rectangles indicated the different clusters. (B) PCA plots based on all SNPs among all individuals. (C) PCA plots among individuals without the outgroup species *P. obovata* (Pab001). (D) A schematic diagram depicting the best-fitting model inferred by fastsimcoal2 with the 95% CI for the estimates of effective population size and event times, assuming a mutation rate µ = 8.0×10^-10^ per base per year and a generation time g = 25 years. The ancestral population is depicted in grey, and the Central-Europe, Finland, Sweden-Norway population is depicted using dark red, dark blue, and dark green, respectively. The inferred demographic parameters are described in the text and shown in Table S4.

The location of samples from southern Sweden and southern Finland in the PCA and the dendrogram relationship suggest that these trees likely have been imported from central Europe sometime during the twentieth century (Chen et al. 2019). We therefore removed these individuals (Pab015 and Pab002) from further analyses. The remaining samples were clustered into three populations corresponding to the groupings seen in the PCA and dendrogram and are hereafter referred to as ‘Central-Europe’, ‘Finland’ and ‘Sweden-Norway’. Earlier research have shown that *P. abies* is subdivided into three main domains, the Alpine domain, the Carpathian domain and the Fennoscandian (Baltico–Nordic) domain (Heuertz et al. 2006; Chen et al. 2019). Our population structure analyses suggest that our Central-Europe population is likely composed of individuals derived from the Carpathian and/or Alpine domains. We also observe sub-structuring (Finland and Sweden-Norway) within the Fennoscandian (Baltico–Nordic) domain that could represent either population structure within the Fennoscandian domain or effects of ongoing hybridisation with *P. obovata* that is more apparent in the eastern (Finland) population (Tsuda et al 2016).

### Low level of genome-wide nucleotide diversity in Norway spruce

Pairwise nucleotide diversity (π) across the *P.abies* genome was relatively low (Table 1). Based on all sites, the Finland population harboured the highest nucleotide diversity with 0.00631±0.00564 (mean ± SD), followed by Sweden-Norway (0.00625±0.00516) with Central-Europe population showing the lowest pairwise nucleotide diversity (0.00494±0.00462) (Table 1). We observed the same pattern across populations when analysing different functional categories of sites separately. One explanation for the higher diversity seen in the Finland and Sweden-Norway populations could be recent material transfers that were introduced for reforestation during the nineteenth and early twentieth century (Chen et al. 2019). Although we removed samples from southern Finland and southern Sweden from our analysis that were likely influenced by materials transfer, there are still possibilities for recent admixture and hybridization between northern and southern individuals given the outbreeding mating system and high levels of wind-dispersed pollen in *P. abies*. Another possible explanation for the higher diversity seen in Finland could also be recent admixture with *P.obovata* (Tsuda et al. 2016). Using nuclear SSR and mitochondrial DNA from 102 and 88 populations, Tsuda et al. (2016) revealed a demographic history where *P.abies* and *P.obovata* have frequently interacted and where migrants originating from the Urals and the West Siberian Plain recolonized northern Russia and Scandinavia using scattered refugial populations of Norway spruce as stepping stones towards the west. This scenario is further supported by exome capture sequencing data which suggest that approximately 17% of *P. abies* in the Nordic domain were derived from *P. obovata* about ∼103K years ago (Chen. et al. 2019).

**Table 1.**
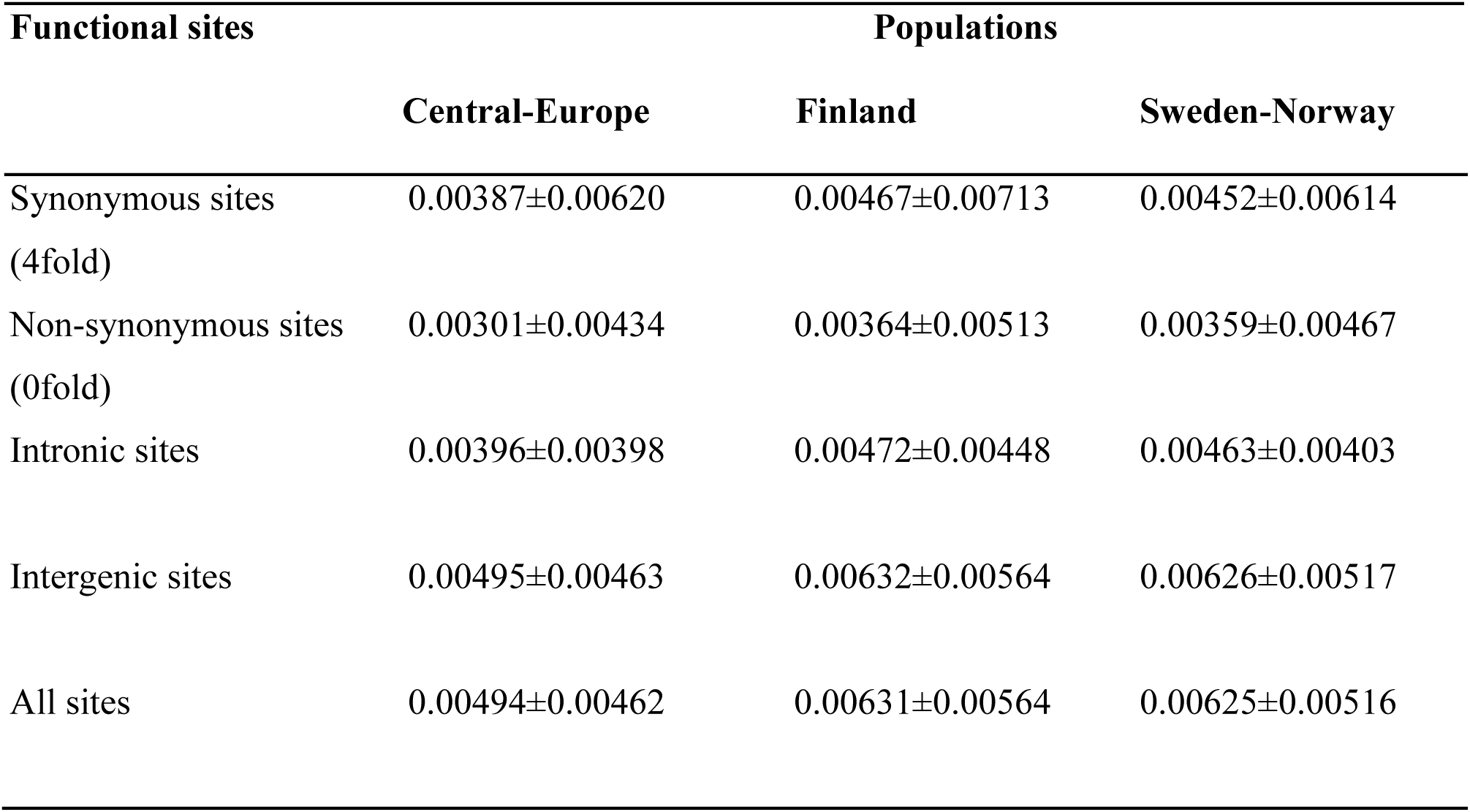
Pairwise nucleotide diversity (π) for different classes of functional sites for three populations of Norway spruce.

For the various genomic contexts we found that levels of nucleotide diversity were highest at intergenic sites, followed by introns, 4-fold synonymous sites with lowest levels of diversity seen at 0-fold non-synonymous sites (Table 1). As every possible mutation at fourfold degenerate sites is synonymous, 4-fold synonymous sites are often assumed to be neutral or nearly neutral (Andolfatto 2005; Filatov et al. 2018) and the lower diversity we observe at non-synonymous sites is thus consistent with purifying selection eliminating deleterious mutations that affect protein coding genes in *P. abies*. Earlier research have also suggested pervasive purifying selection in *P. abies* by showing lower frequency of differences at non-synonymous SNPs relative to silent sites based on pooled population sequencing (Chen et al. 2016). In contrast to variants located in genes, intergenic regions had substantially higher levels of nucleotide diversity in all three populations, which may reflect either relaxed selective constraints or possibly higher mutation rates in these regions. Another possible explanation is that read mapping errors are higher in intergenic regions since these regions contain a greater fraction of repetitive sequences. We have tried to filter away such regions (see Material and Methods) and we do not observe a strong correlation between intergenic diversity and repeat density (Supplementary Figure S6). However, we cannot be entirely sure that we have successfully eliminated all such regions. Higher nucleotide diversities in intergenic regions have been observed also in the chloroplast genome of *P. abies* (Sullivan et al. 2017), as well as in the nuclear genome of other plants, such as *Populus* (Wang et al. 2016), or animals (collared flycatchers, Dutoit et al. 2017).

Earlier studies of nucleotide diversity in *P.abies* have mainly targeted coding regions. For instance, Heuertz et al. (2006) found a mean nucleotide diversity of 0.0021 across all sites based on data from 22 loci analysed in 47 Norway spruce samples collected from seven natural populations. Larsson et al. (2013) found a mean nucleotide diversity (π) of 0.0047 (SD=0.0034) based on 11 loci screened in ten natural *P. abies* populations from across Scandinavia. More recently, Chen et al. (2019) used exome capture data in *P. abies* to show that nucleotide diversity at 4-fold synonymous sites ranged from 0.0072 to 0.0079 in the three domains while 0-fold nucleotide diversity ranged from 0.0027 to 0.0032. Our results, based on whole genome data, produce slightly different estimates of nucleotide diversity but are of the same order of magnitude as earlier estimates. The reasons for the slight discrepancies between the different studies are likely explained by differences in sampling design and sequencing strategies.

Compared to other species, estimates of nucleotide diversity in Norway spruce (0.0049-0.0063) are slightly higher than that in the predominantly autogamous legume *Medicago truncatula* (π=0.0043, Branca et al. 2011), but lower than in maize (π=0.0066, *Zea mays* L.) (Gore et al. 2009), wild rice (π=0.0077, *Oryza rufipogon* and *Oryza nivara)* (Xu et al. 2012), and three to five-fold lower than estimates in *Populus* (π=0.0133, *Populus. tremula*; π=0.0144, *P. tremuloides*) (Wang et al. 2016). Our results thus indicate that Norway spruce is characterized by a relatively low level of nucleotide diversity, especially when factoring in the extensive distribution range of the species. Norway spruce (*P. abies*) is one of the dominant tree species in the boreal and temperate zones of Europe, with a wide geographical distribution. *P. abies* has a longitudinal range ranging from the French Alps (5°27′E) to the Ural mountains (59°E), a latitudinal range from the southern Alps to Northern Scandinavia (70°N) and an altitudinal range from sea level to well above 2,300 m in the Italian Alps (Jansson et al. 2013). *P. abies* has also been planted widely outside its natural range and currently plays a major role in European forestry. Our results thus illustrate a disparity between the extensive geographic distribution of Norway spruce and an overall low level of genome-wide genetic diversity.

### Divergence and population fluctuations during the demographic history of *P. abies*

One possible explanation for the low levels of genetic diversity seen in Norway spruce is demographic history. If *P. abies* has experienced sharp reductions in the effective population size due to environmental events e.g. climate change, in the past and have only recently become abundant again, observed levels of genetic diversity would be substantially lower than expected based on current distribution and population sizes.

To assess whether demographic factors are responsible for the low genetic diversity seen in *P.abies*, we inferred the demographic history for all populations using the joint site frequency spectrum (SFS) based on all SNPs using coalescent simulations implemented in fastsimcoal2 (Excoffier et al. 2013). A mutation rate of 8.0×10^-10^ per site per year and a generation time of 25 years (De La Torre et al. 2017; Chen et al. 2019) were assumed to calculate the parameter estimates of the divergence times and effective population sizes, and their associated 95% confidence intervals (CIs) based on 100 parametric bootstraps. We evaluated eight demographic models that consider a range of scenarios for effective population size changes (Supplementary Figure S3). The best-fitting model was model ‘Pop012-Bot’ which was chosen based on Akaike’s weight of evidence ≈ 1 (Supplementary Table S3). In this model, a common ancestral population experience a population contraction followed by the split of the Central-Europe population, followed by the divergence of the Sweden-Norway and Finland populations. All three sub-populations subsequently experience bottlenecks that lasted until ∼ 28,5 kya (1.65-103.1 kya) (Figure 1D; Supplementary TableS4). The ancestral effective population size of *P. abies* was much larger than current day populations, with an estimated effective population size of 76,515-457,249 (95% CI). This population went through a sharp bottleneck, reducing the effective size to 2,216-24,608 approximately 45.6 million years ago (mya) (12.7-62.7 mya). This bottleneck may reflect the influence from the Paleogene, when the global climate showed a significant cooling trend in the beginning of the Oligocene epoch (Vandenberghe et al. 2012). The divergence time for the Central-Europe population is estimated to have occurred about 114.4 kya (111.2-119,8 kya), whereas the divergence between the Finnish and Swedish-Norwegian population is more recent, dated to around ∼ 32.7 kya (28.7-44.2 kya). All three populations went through subsequent bottlenecks (N_e_ in Central-Europe ∼ 209-5,011; N_e_ in Sweden-Norway ∼ 1,429-44,282; Ne in Finland ∼179-41,190), which timing roughly corresponds to when several abrupt changes in temperature took place during the quaternary, including the Last Glacial Maximun (LGM).

The inferred demographic history for the Central-Europe population, which appears to have experienced a longer period of the most recent bottleneck, coincides with this population also having the lowest levels of nucleotide diversity. Mountain regions are known to be habitats suitable for species survival during drastic climate fluctuations since they allow species to respond to climatic change by moving along a altitudinal gradient and such movements are further expected to promote population divergence on a regional scale (Tollefsrud et al. 2008). Southern areas in Europe, such as the Alps and Carpathians, as well as the mountains adjacent to the northern Scandinavian coastline have already been identified as likely refuges for many species during the last glacial age (Dahl 1995; Tollefsrud et al. 2008; Parducci et al.2012; Arukwe and Langeland 2013), lending credibility to the patterns of population divergence and size changes we observed in our data. Recent bottlenecks for different populations of *P. abies* have also been identified in earlier studies (Heuertz et al. 2006; Chen et al. 2019). Another reason that may help to explain the low levels of nucleotide diversity we observe in *P. abies* is that most regions were likely already forested before the postglacial arrival of *P. abies*, making colonizing populations more prone to genetic drift induced by founder events or subsequent bottlenecks that act to reduce genetic diversity (Tollefsrud et al. 2008). The demographic history of *P. abies* is thus an important factor explaining the low levels of nucleotide diversity observed. However, the limited sampling scheme employed in this study, together with the uncertain estimates of mutation rates and generation times suggest that the exact timing and size of population changes should be treated with a healthy amount of scepticism.

### Pervasive molecular signal of natural selection by linkage

Genetic hitchhiking is the process by which an allele experiences a change in frequency because it is physically located in the vicinity of another allele that is under positive selection. Such selective sweeps will consequently reduce the level of diversity not only at the selected locus but also at nearby sites (Smith and Haigh 1974). Similarly, background selection also removes genetic diversity at sites linked to deleterious mutations that are under purifying selection in a population (Charlesworth et al. 1995). Therefore, natural selection via either positive selection favouring advantage mutations (“hitchhiking”) or purifying selection against deleterious mutations (“background selection”) is expected to also affect standing levels of genetic diversity in a species (Corbett-Detig et al 2015).

If natural selection is widespread across the genome, it can lead to a reduction in neutral genetic diversity in regions of low recombination (Begun and Aquadro 1992) since sites in such regions are expected to experience stronger effects of linked selection. Such patterns have already been observed across a wide range of animals and plants, and has been used to support the view that linked selection is a powerful force affecting genome-wide level of genetic diversity in many species (Corbett-Detig et al. 2015). In accordance with the expectations from linked selection, we observe a positive correlation between nucleotide diversity at 4-fold synonymous sites and an LD-based estimate of the population-scaled recombination rate in all three populations. A simple linear regression model further showed that variation in recombination rate explained 2.6-3.4% of the genome-wide levels of variation at neutral 4-fold synonymous sites in the three populations (Figure 2). We observed the weakest association between neutral 4-fold diversity and recombination rate in the Central-Europe population, likely reflecting the more severe and extended bottleneck seen in this population (Figure 1D). When we analyse correlations between recombination rates and nucleotide diversity at 0-fold non-synonymous sites, sites located in introns or in intergenic regions, we also observe positive correlations. Across all genomic contexts, the strongest correlation between diversity and recombination rate was observed at intergenic sites, ranging from 0.419 to 0.601 in the three populations (Figure 2). In line with this, a relatively large fraction (22.2% - 26.0%) of the variation in genetic diversity can be explained by variation in the recombination rate at intergenic sites (Figure 2). The higher levels of nucleotide diversity we observe in intergenic regions can at least partly explain this, as the power to detect any relationship between diversity and recombination rate is likely also higher in these regions (Table 1).

**Figure 2.**
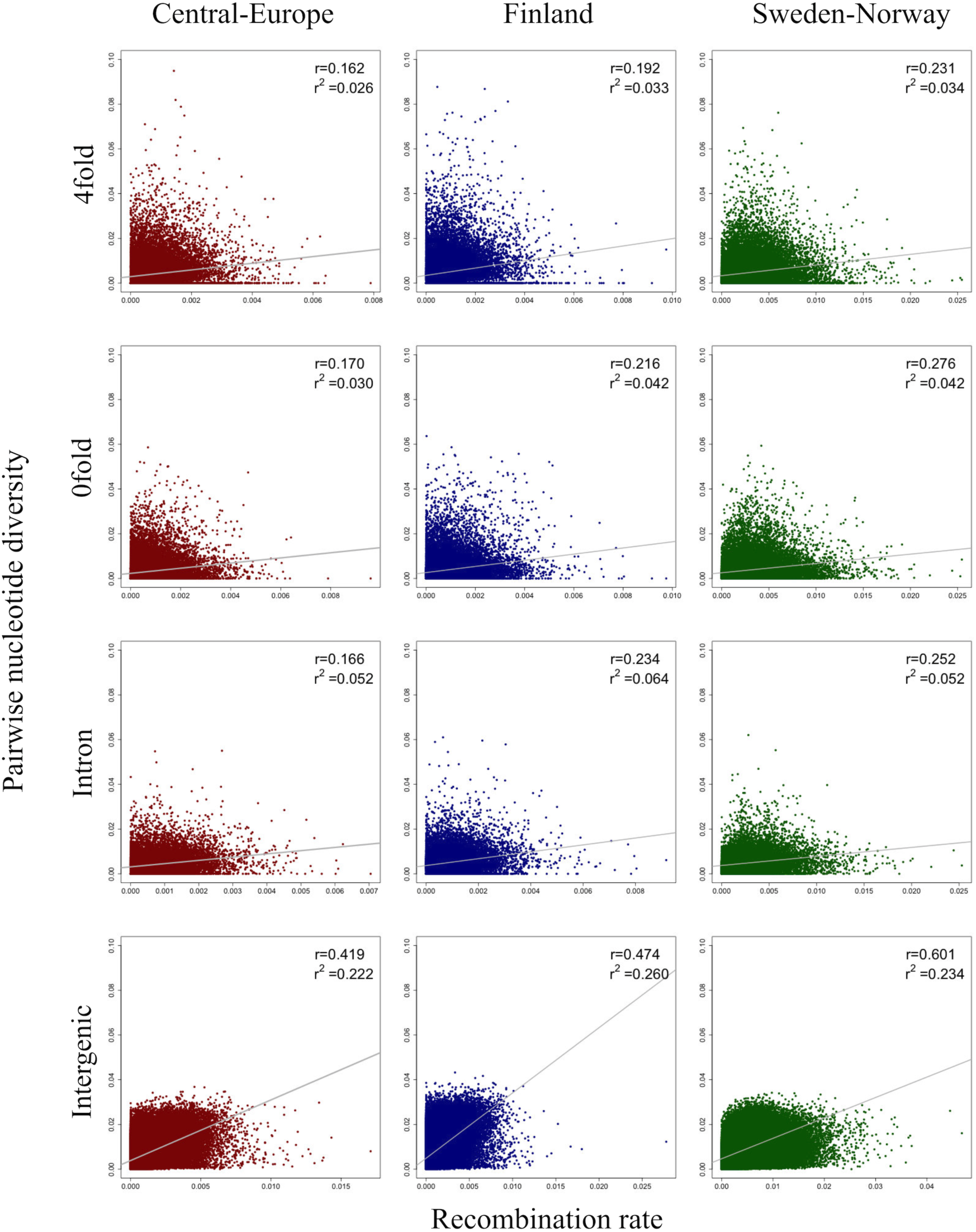
Scatterplots of 4-fold, 0-fold, intronic and intergenic nucleotide diversity versus the population-scaled recombination rates across the *P. abies* genome for Central-Europe (dark red), Finland (dark blue), and Sweden-Norway (dark green). The Spearman correlation coefficient (r) and linear regression (r^2^) are shown in the top right of each plot. Best fitting regression lines are depicted in grey.

If the positive relationship between nucleotide diversity and recombination rate was merely caused by the mutagenic effect of recombination, similar patterns should also be observed between divergence and recombination rate (Kulathinal et al. 2008; Wang et al. 2016). We therefore also assessed correlations between divergence and recombination rate at the different classes of sites (4-fold, 0-fold, introns and intergenic sites) and the results show no correlations (linear regression r^2^ <1%, Figure 3). The positive correlation we observe between nucleotide diversity and recombination rate is consequently driven by linked selection rather than through mutagenic recombination.

**Figure 3.**
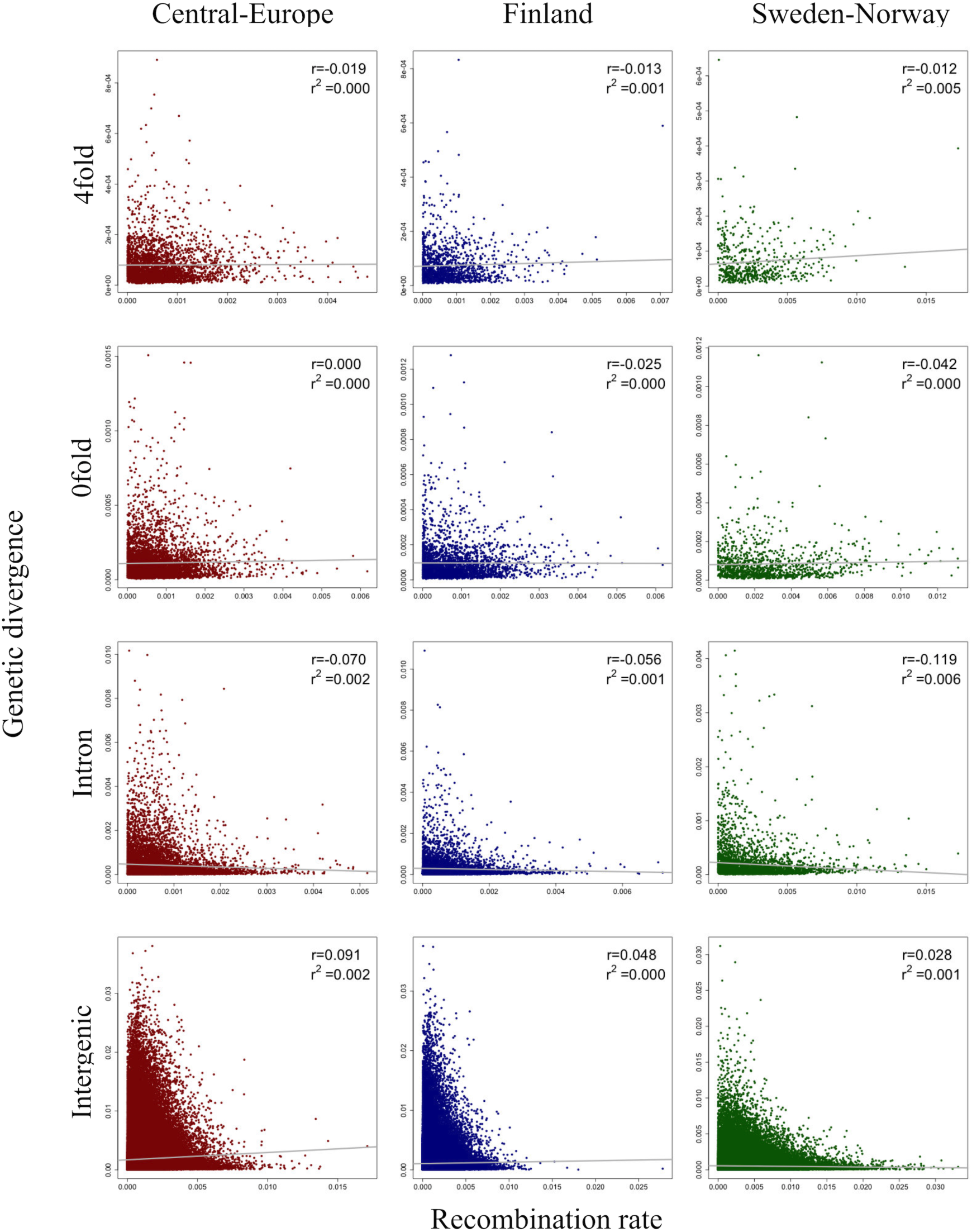
Scatterplots of 4-fold, 0-fold, intronic and intergenic genetic divergence versus the population-scaled recombination rates across the *P. abies* genome for Central-Europe (dark red), Finland (dark blue), and Sweden-Norway (dark green). The Spearman correlation coefficient (r) and linear regression (r^2^) are shown in the top right of each plot. Best fitting regression lines are depicted in grey.

Levels of nucleotide variation is expected to be proportional to the density of selected sites per genetic map unit (Flowers et al. 2011), since regions with a higher density of mutations under selection will experience stronger effects of linked selection. If genic regions represent the most likely targets of natural selection, we consequently expected to see a negative association between the density of genic sites and levels of nucleotide diversity. This observation may help explain the negative correlation between polymorphism and gene density seen in many plant species, e.g. *Arabidopsis thaliana* (Nordborg et al. 2005), Asian rice (*Oryza sativa* ssp. *Japonica*; ssp. *indica*; *O. rufipogon*) (Flowers et al. 2011) or insects (eg. *Heliconius melpomene*, Martin et al. 2016). Gene density is normally scored as the number of protein-coding genes in a region of fixed size along a chromosome. However, due to the fragmented nature of the *P. abies* genome, we are unable to assess gene density in this manner. Instead, to assess the effects of the density of putative sites under selection, we measured the density of genic sites on a per scaffold basis, estimated as the number of base pairs that were annotated as coding over the length of the scaffold. We observed a negative association between our measure of coding density and nucleotide diversity both at 0-fold non-synonymous, 4-fold synonymous and intergenic sites (Figure 4; Supplementary Figure S4). We found no correlation between gene density and diversity at intronic sites and the impact of gene density on diversity in introns was negligible (<1%) (Supplementary Figure S4). The strongest correlation we observed was between gene density and diversity of 0-fold sites, possibly reflecting that non-synonymous substitutions are under stronger selection. However, spurious results can be generated when two explanatory variables co-vary but are analysed separately, for example, a negative covariation of recombination rate and gene density will strengthen the signatures of linked selection across the genome (Flowers et al. 2012; Ellegren and Galtier 2016). We therefore also assessed the correlation between recombination rate and gene density. We found no correlation between gene density and recombination rates in Central-Europe or Finland population (linear regression r^2^ <1%) (Supplementary Figure S5). However, we detect a significantly negative correlation in the Sweden-Norway population (Figure S5), which could help explain why we also observed the strongest negative correlation between functional diversity and gene density in this population. We did not observe correlations (Supplementary Figure S6; linear regression r^2^ <1%, data not shown) between nucleotide diversity and GC density or repeat density, measured as the fraction of sites per scaffold that are annotated as repeats.

**Figure 4.**
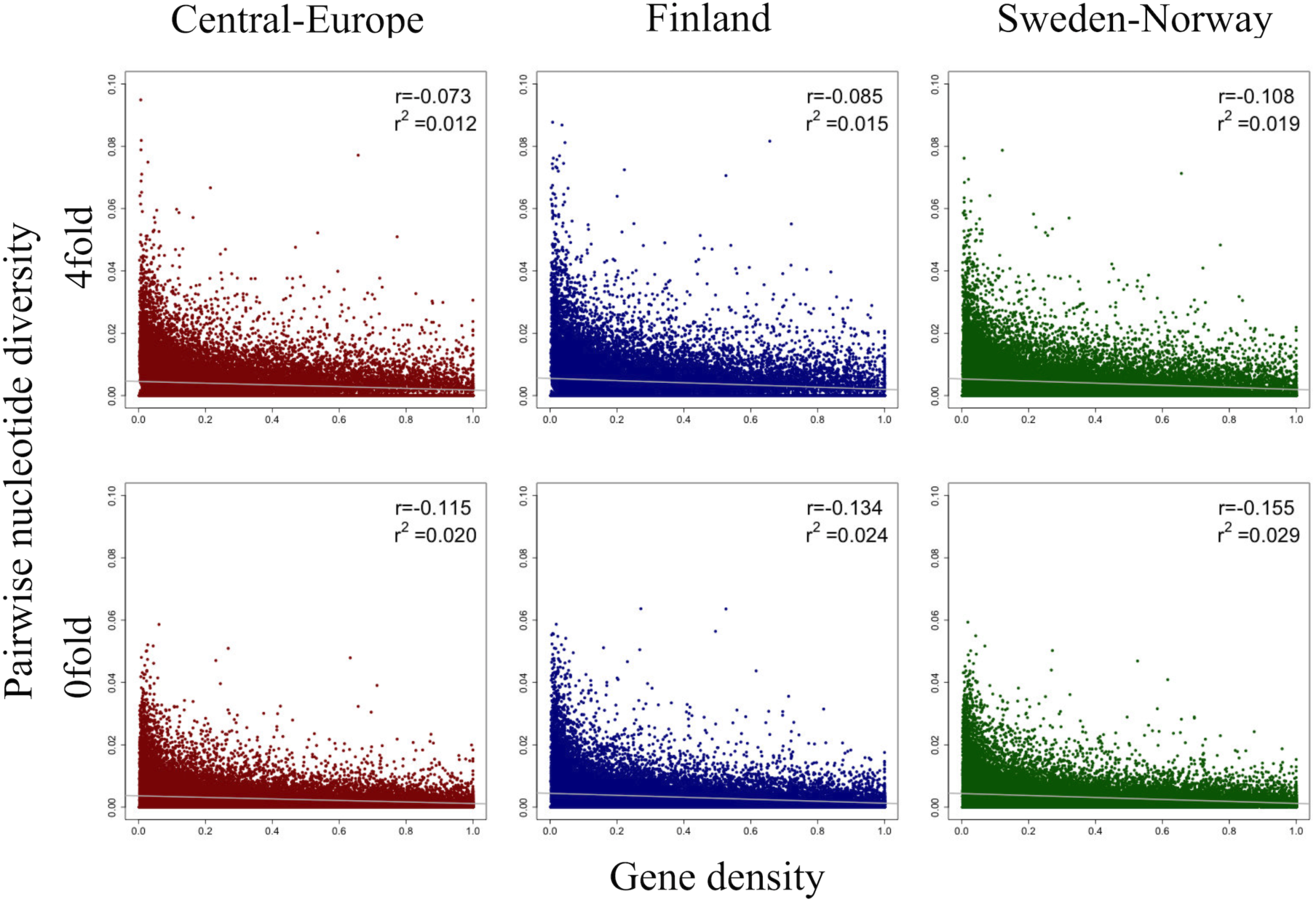
Scatterplots of 4-fold and 0-fold genetic diversity versus gene density for Central-Europe (dark red), Finland (dark blue), and Sweden-Norway (dark green). The Spearman correlation coefficient (r) and linear regression (r^2^) are shown in the top right of each plot. Best fitting regression lines are depicted in grey.

Overall, our results suggest that widespread natural selection contributes to variation in the genome-wide levels of nucleotide diversity in *P. abies*. Similar effects have been found in a wide variety of species, e.g. *Drosophila melanogaster* (Begun and Aquadro 1992), humans (Hellman et al. 2003), *Caenorhabditis elegans* (Cutter and Payseur 2003), *Saccharomyces cerevisiae (*Cutter and Moses 2011*), Populus* (Wang et al. 2016) and *Emiliania huxleyi* (Filatov et al. 2018). We identified positive correlations between diversity and recombination rate for both genic region and intergenic region in all three populations, but no correlations between divergence and recombination rate, indicating pervasive effects of linked selection across the *P. abies* genome. Purifying selection against deleterious mutations in regions of higher gene density is more likely to account for the negative relationship between gene density and neutral genetic diversity, and the magnitude of such effects depends on the strength of purifying selection (Sella et al. 2009). The weak negative correlations we observed between nucleotide diversity (0-fold, 4-fold and intergenic region) and gene density suggest that background selection may be limited in *P. abies*.

Finally, natural selection is expected favour lower mutation rates because most spontaneous mutations are deleterious, and thus species with larger population sizes tend to have lower mutation rates because larger populations experience stronger effects of natural selection (Lynch et al. 2016; Xu et al. 2019). Therefore, a low mutation rate could at least partly explain low levels of nucleotide diversity in species with extremely large census population sizes. For instance, Xu et al. (2019) used re-sequencing from 68 samples spanning the global distribution of giant duckweed (*Spirodela polyrhiza*) to show that this species has extremely low intraspecific genetic diversity (e.g. π_synonymous_=0.00093) despite achieving extremely high census population sizes in nature due to high rates of clonal growth. They also estimated a very low spontaneous mutation rate (at least seven times lower than estimates in other multicellular eukaryotes). This is an example of a species showing a low level of genetic diversity despite having a wide current natural distribution and where selection may have reduced mutation rates to very low levels. De La Torre et al. (2017) showed that the mutation rate of gymnosperms are, on average, seven times lower than in angiosperms based on data from a set of 42 orthologous, single-copy genes. A low mutation rate might thus also help explain the low levels of nucleotide diversity that are observed across most conifers (see Introduction).

## Conclusion

Under the neutral theory species with larger effective population size are expected to have higher levels of genetic diversity (Kimura 1983). However, observations have suggested that the range of effective population sizes across species exceeds the observed range of genetic diversity by several orders of magnitude (Lewontin 1974; Leffler et al. 2012) and this long-standing issue is now known as Lewontin’s paradox (Lewontin 1974). Norway spruce has a wide natural geographic distribution across the northern hemisphere (Farjon 1990; Jansson et al. 2013) and is often the dominant tree species in many boreal forests. Previous genomic studies have suggested a relatively low level of nucleotide diversity (e.g. 0.0021 in Heuertz et al. 2006, 0.0047 in Larsson et al. 2013) compared to many angiosperm species with similar distribution ranges. Our analyses of nucleotide diversity in *P. abies* by whole genome re-sequencing data support the view that genome-wide levels of nucleotide diversity is low and of the same order of magnitude as seen in earlier studies, even if these have been based almost exclusively on a limited set of short genomic fragments from predominantly coding regions.

We have focused on identifying potential factors that can help explain the low levels of genetic diversity seen in *P. abies*. Population structure in our sample is characterised by three relatively well-defined clusters that closely correspond to the geographic origin of samples from Central Europe, Finland, Sweden/Norway. The demographic history for all three populations is characterised by an ancient bottleneck in an ancestral population and more recent bottlenecks that coincides with the last glacial maximum across all populations. Population size has only recently recovered, suggesting that reductions in effective population size throughout the history of the species is an important contributor to the low intraspecific diversity. We also observe a positive correlation between nucleotide diversity and recombination rate across both genic and intergenic regions, suggesting widespread action of linkage selection across the *P. abies* genome and such linked selection can also contribute to the species-wide loss of genetic diversity. A weak, but significant negative correlation between nucleotide diversity (at 0-fold, 4-fold and intergenic sites) and gene density is also observed, which could be interpreted as background selection only having weak effects in Norway spruce. Variation in linked selection across the *P. abies* genome is thus also a likely contributing factor explaining the low genetic diversity in the species. Finally, the unusually low mutation rate, common to most conifers, is yet another factor that contributes to the low species-wide diversity in *P. abies*.

Our results provide the first insights into whole-genome levels of variation in a conifer species. This allows us to also assess patterns of variation with a substantially higher resolution than earlier studies and lays the foundation for more accurate inferences in *P. abies*. Our results suggest that both past demographic events as well as pervasive effects of linked selection have important influences on genome-wide levels of diversity. Earlier studies have suggested that demographic events could result in the detection of spurious signatures of natural selection (Fay et al. 2001) while the presence of linked selection could also bias inferences of demographic history (Slotte 2014). Thus new models that allow for the joint estimation of demography and selection from genome-wide data are urgently needed to assess the relative importance of demography and linked selection for explaining patterns of genome-wide variation in genetic diversity in *P. abies*.

## Acknowledgements

The research has been funded by grants from the Knut and Alice Wallenberg foundation (Norway spruce genome project) and the Swedish Foundation for Strategic Research (SSF, grant number RBP14–0040). Data generation was supported by Science for Life Laboratory and the National Genomics Infrastructure (NGI) which provided access to massive parallel sequencing. All analyses were performed on resources provided by the Swedish National Infrastructure for Computing (SNIC) at Uppsala Multidisciplinary Centre for Advanced Computational Science (UPPMAX) under the projects b2012141, SNIC 2017/1-438, SNIC 2018/3-529 and uppstore2017066.

