## Supplementary Figures and Tables for "Demography and natural selection have shaped genome-wide variation in the widely distributed conifer Norway Spruce (*Picea abies*)"

Table S1. Summary statistics of Illumina re-sequencing data per sample.

| Sample ID | Location | Sequence Platform | Sequence Coverage (x) | BAM Coverage (x) |
|---|---|---|---|---|
| Pab001[a] | Yakutsk, Russia | HiSeq 2000 | 10.1 | 12.6 |
| Pab002 | Gettinge, Sweden | HiSeq 2000 | 36.9** | 12.0 |
| Pab003 | Vitebsk, Belarus | HiSeq 2000 | 9.5 | 12.2 |
| Pab004 | Blizyn, Poland | HiSeq 2000 | 10.5 | 11.8 |
| Pab005 | Toplita, Romania | HiSeq 2000 | 10.3 | 12.2 |
| Pab006* | Köttsjön, Sweden | HiSeq 2000 | 50.0** | 11.1 |
| Pab007 | Hatfjelldal, Norway | HiSeq X | 12.6 | 14.5 |
| Pab008 | Rovaniemen, Finland | HiSeq X | 11.8 | 14.5 |
| Pab009 | Kittilä, Finland | HiSeq X | 13.2 | 14.2 |
| Pab010 | Suomussalmi, Finland | HiSeq X | 11.9 | 14.5 |
| Pab011 | Hemnes, Norway | HiSeq X | 14.6 | 14.2 |
| Pab012 | Levanger, Norway | HiSeq X | 12.4 | 14.5 |
| Pab013 | Grane, Norway | HiSeq X | 13.3 | 15.1 |
| Pab014 | Tyda, Norway | HiSeq X | 12.4 | 15.2 |
| Pab015 | Loppi, Finland | HiSeq X | 13.6 | 14.9 |
| Pab016 | Marsfjället, Sweden | HiSeq X | 19.6 | 15.5 |
| Pab017 | Marsfjället, Sweden | HiSeq X | 21.0 | 15.5 |
| Pab018 | Marsfjället, Sweden | HiSeq X | 20.4 | 15.8 |
| Pab019 | Marsfjället, Sweden | HiSeq X | 20.2 | 25.5 |
| Pab020 | Marsfjället, Sweden | HiSeq X | 21.1 | 25.2 |
| Pab021 | Marsfjället, Sweden | HiSeq X | 20.9 | 25.5 |
| Pab022 | Marsfjället, Sweden | HiSeq X | 19.9 | 26.0 |
| Pab023 | Marsfjället, Sweden | HiSeq X | 19.8 | 25.2 |
| Pab024 | Marsfjället, Sweden | HiSeq X | 18.9 | 24.5 |
| Pab025 | Marsfjället, Sweden | HiSeq X | 18.8 | 23.8 |
| Pab026 | Långrumpskogen, Sweden | HiSeq X | 19.2 | 23.8 |
| Pab027 | Långrumpskogen, Sweden | HiSeq X | 18.6 | 23.5 |
| Pab028 | Långrumpskogen, Sweden | HiSeq X | 19.3 | 23.5 |
| Pab029 | Långrumpskogen, Sweden | HiSeq X | 18.5 | 23.2 |
| Pab030 | Långrumpskogen, Sweden | HiSeq X | 18.0 | 23.2 |
| Pab031 | Långrumpskogen, Sweden | HiSeq X | 17.8 | 23.2 |
| Pab032 | Långrumpskogen, Sweden | HiSeq X | 20.1 | 23.5 |
| Pab033 | Långrumpskogen, Sweden | HiSeq X | 20.0 | 23.6 |
| Pab034 | Långrumpskogen, Sweden | HiSeq X | 20.2 | 23.1 |
| Pab035 | Långrumpskogen, Sweden | HiSeq X | 19.3 | 23.3 |
| Mean | | | 18.13 | 18.85 |

[a] *Picea obovata.*

*reference individual Z4006 (Nystedt et al. 2013).

** Down-sampled to match remaining samples.

Table S2. Summary statistics of genomic subsets with each included 35 individuals. Genomic length (mega bases) is the length of combined scaffolds in each subset; Average scaffold length (kilo bases) is the average length of each scaffold included in the subset; Unfiltered recodes (in millions) and unfiltered SNPs (in millions) are number of recodes and number of SNPs present in the unfiltered raw VCF files, respectively; Hard filtered SNPs (in millions) are number of SNPs retained after performing hard filtering criteria, and hard filtered scaffolds (%) are percentage of the scaffolds that contained only filtered SNPs in each subset.

| Subset | Genomic Length (Mb) | Average Scaffold length (KB) | Unfiltered Recodes (Million) | Unfiltered SNPs (Million) | Hard filterec SNPs (Million) | Hard filtered Scaffolds (%) |
|---|---|---|---|---|---|---|
| 1 | 2,654.8 | 26.6 | 230.7 | 217.7 | 114.1 | 96.1 |
| 2 | 1,657.9 | 16.6 | 113.2 | 107.8 | 44.1 | 80.1 |
| 3 | 451.9 | 4.5 | 40.6 | 38.4 | 15.1 | 77.6 |
| 4 | 394.3 | 3.9 | 35.4 | 33.4 | 12.9 | 76.2 |
| 5 | 481.1 | 4.8 | 43.9 | 41.4 | 16.4 | 79.4 |
| 6 | 243.3 | 2.4 | 21.7 | 20.5 | 7.6 | 70.4 |
| 7 | 245.3 | 2.5 | 22.0 | 20.7 | 7.7 | 70.6 |
| 8 | 257.8 | 2.6 | 23.2 | 21.9 | 8.4 | 71.9 |
| 9 | 329.2 | 3.3 | 27.5 | 25.8 | 11.3 | 64.7 |
| 10 | 256.4 | 2.8 | 7.8 | 7.6 | 0.5 | 16.5 |
| 11 | 241.0 | 2.4 | 13.0 | 12.6 | 1.5 | 33.8 |
| 12 | 159.1 | 1.6 | 13.3 | 12.6 | 3.9 | 55.5 |
| 13 | 179.2 | 1.8 | 15.0 | 14.2 | 4.5 | 56.6 |
| 14 | 196.9 | 2.0 | 16.5 | 15.7 | 5.3 | 57.9 |
| 15 | 213.0 | 2.1 | 17.9 | 17.0 | 6.0 | 58.5 |
| 16 | 229.6 | 2.3 | 19.2 | 18.2 | 6.6 | 59.7 |
| 17 | 237.6 | 2.4 | 17.6 | 16.8 | 5.2 | 59.3 |
| 18 | 262.2 | 2.6 | 20.1 | 19.2 | 6.1 | 66.3 |
| 19 | 331.3 | 3.3 | 25.7 | 24.4 | 8.4 | 66.1 |
| 20 | 433.6 | 6.2 | 25.3 | 23.6 | 8.2 | 39.5 |
| Total | 9,455.5 | 4.8 | 749.6 | 709.5 | 293.9 | 63.2 |

Table S3. Relative likelihood of different models.

| Model | Max ($log_{10}$($Lhood_i$))[a] | No. of parameters (d) | $AIC_i$[b] | $\Delta_i$ [b] | Model normalized relative likelihood (w )[b] |
|---|---|---|---|---|---|
| No-Bot | -433738283 | 14 | 1997798560 | 1755486 | ~0 |
| Pop0-Bot | -433378378 | 22 | 1996140853 | 97779 | ~0 |
| Pop1-Bot | -433379115 | 22 | 1996144248 | 101174 | ~0 |
| Pop2-Bot | -433477940 | 22 | 1996599436 | 556362 | ~0 |
| Pop01-Bot | -433373421 | 28 | 1996118033 | 74959 | ~0 |
| Pop02-Bot | -433413182 | 28 | 1996301172 | 258098 | ~0 |
| Pop12-Bot | -433376659 | 26 | 1996132943 | 89869 | ~0 |
| Pop012-Bot | -433357145 | 32 | 1996043074 | 0 | 1 |

[a] Based on the best likelihood among the 50 independent runs for each model.

[b] The calculation of $AIC_i$, $\Delta_i$ and $w_i$ are according to the methods shown in Excoffier et al. (2013).

Table S4. Inferred parameters of demographic history in *P. abies* under best model-Pop012-Bot.

| Parameters | Point estimation | 95% CI[a] | |
|---|---|---|---|
| | | Lower bound | Upper bound |
| NANCTHR | 194773 | 76515 | 457249 |
| NANCSHR[b] | 27223 | 2216 | 24608 |
| NANCTWO | 20211 | 616 | 327809 |
| NPOPANC0[b] | 60478 | 4325 | 47225 |
| NPOPBOT0 | 3393 | 209 | 5011 |
| NPOP0 | 311241 | 78122 | 424731 |
| NPOPANC1 | 92473 | 4041 | 453545 |
| NPOPBOT1 | 14992 | 179 | 41190 |
| NPOP1 | 1633 | 1036 | 376945 |
| NPOPANC2 | 227495 | 11658 | 496858 |
| NPOPBOT2 | 17178 | 1429 | 44282 |
| NPOP2 | 4428 | 2482 | 6489 |
| TBOT012 | 28463 | 1650 | 103125 |
| TENDBOT12 | 30181 | 23788 | 39806 |
| TDIV1 | 32656 | 28669 | 44206 |
| TENDBOT0 | 101338 | 31144 | 109656 |
| TDIV2 | 114400 | 111238 | 119831 |
| TSHREND | 45632194 | 12720263 | 62706394 |

[a] Parametric bootstrap estimates obtained by parameter estimation from 100 datasets simulated according to the overall maximum composite likelihood estimates shown in point estimation columns. Estimations were obtained from 100,000 simulations per likelihood.

[b] Two parameters were very ancient effective population sizes, so that point estimations were not inferred accurately and thus out of corresponding 95% CI ranges.

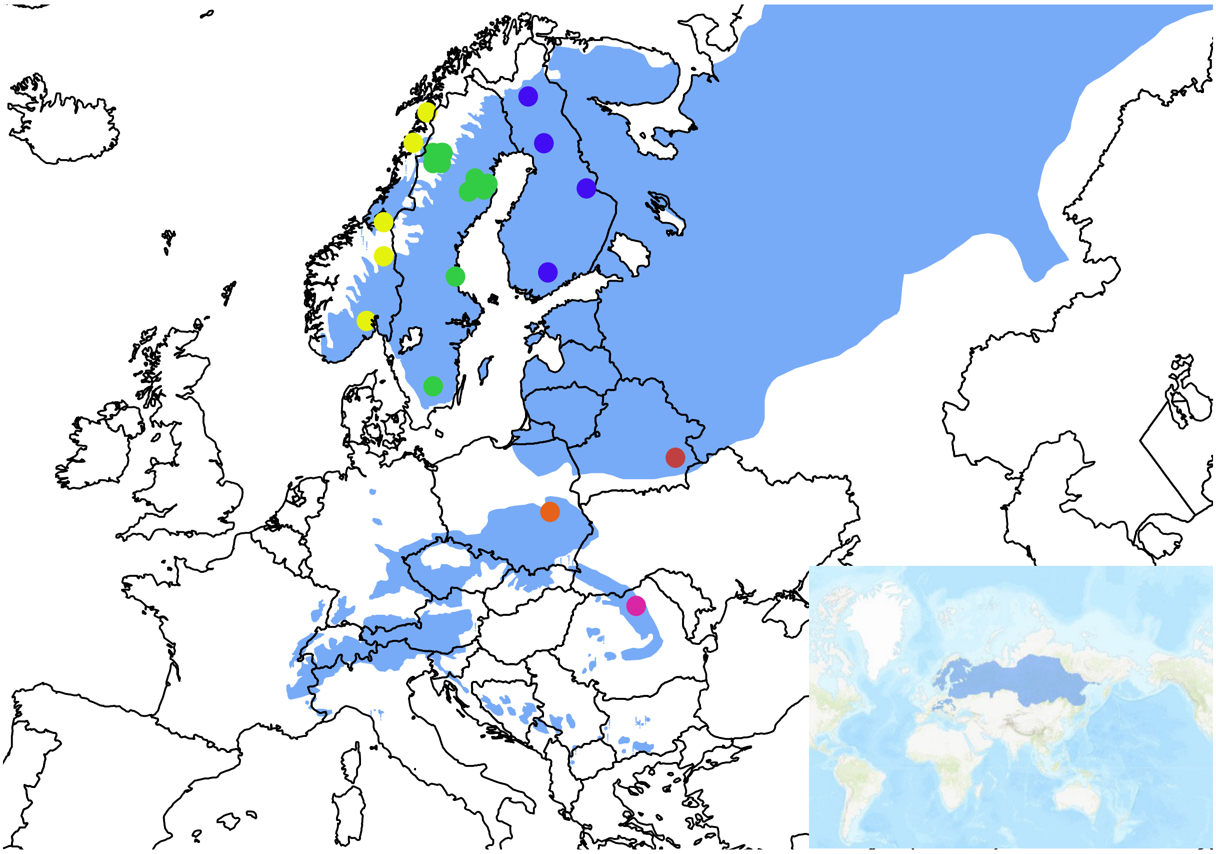

Fig. S1. Geographic distribution of whole-genome re-sequenced individuals of *P. abies*. Individuals from Norway (yellow), Sweden (green), Finland (blue), Poland (orange), Belarus (red), and Romania (pink) are shown in circles with different colors. Individual Pab001 of *P. obovata* located in Yakutsk, Russia is not shown here. The bottom right figure represents the natural distribution of *P. abies* with dark blue color (EUFORGEN 2009,www.euforgen.org).

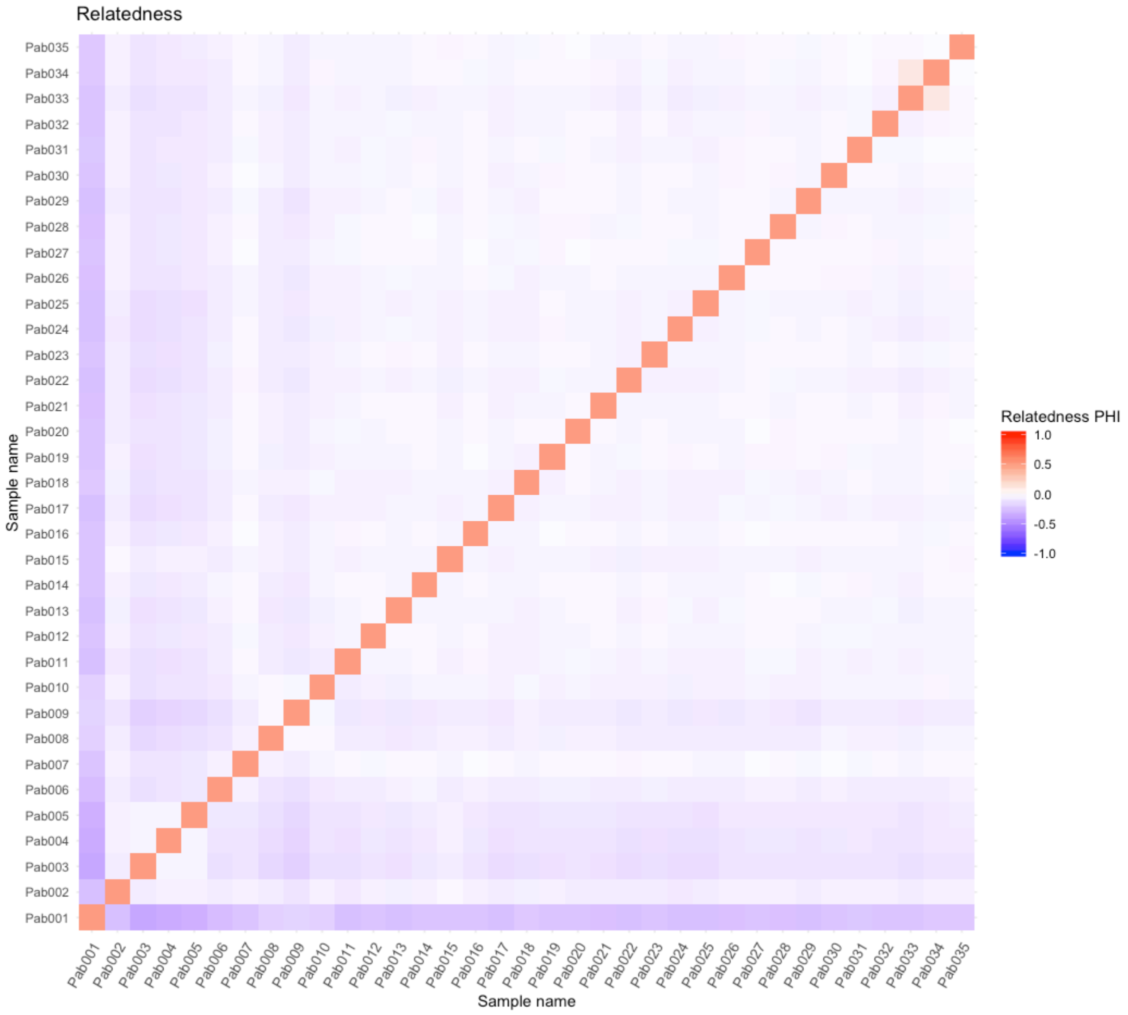


Fig. S2. Estimated genetic relatedness between each pair of 35 individuals using SNP data.

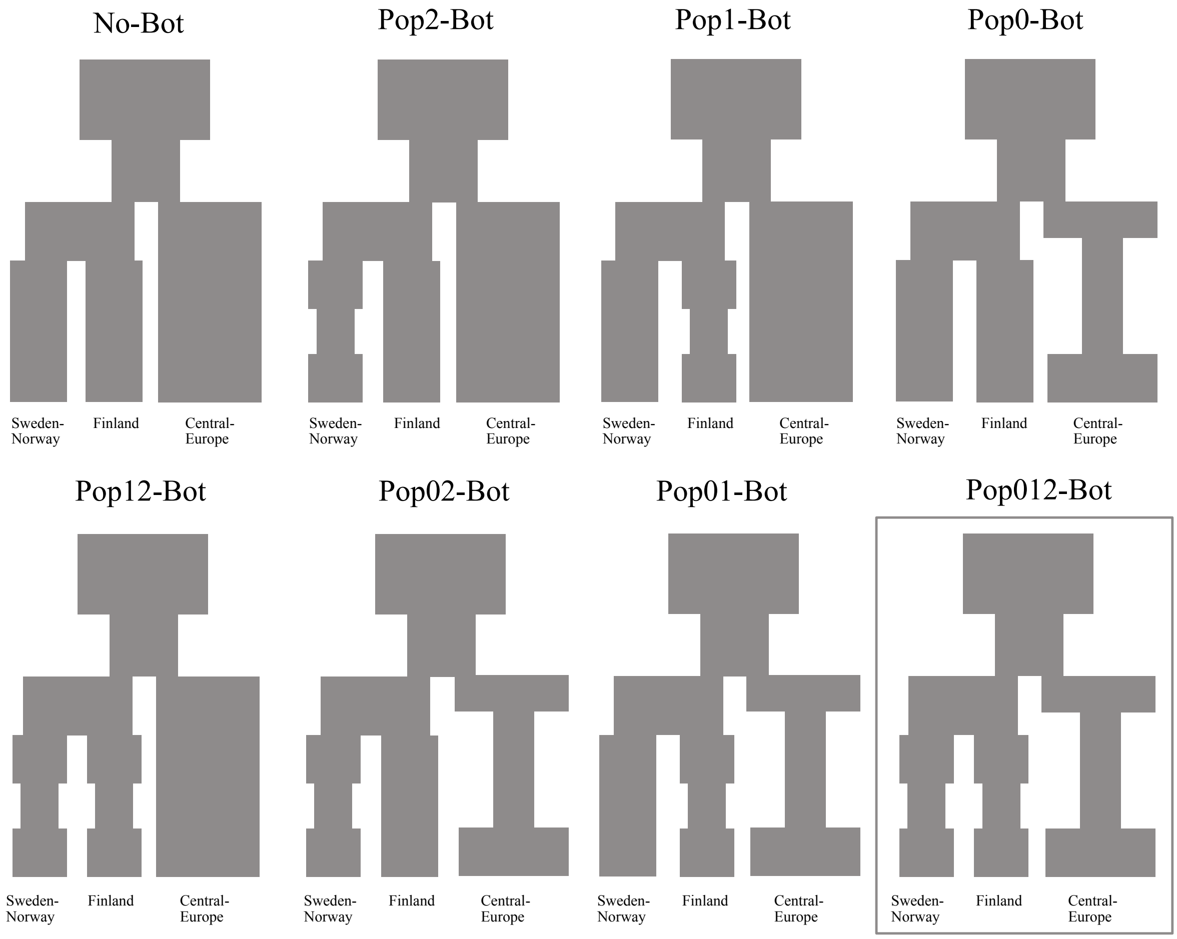


Fig. S3. Schematic diagram of tested demographic models used in fastsimcoal2. All models included three populations and began with common ancestral population experiencing a population bottleneck, followed by first split of the Central-Europe population, followed by the divergence between of the Finland and Sweden-Norway populations. These models differed depending on population sizes after divergence and whether individual populations went through further bottlenecks. Pop0, pop1, pop2 represent Center-Europe, Finland and Sweden-Norway population, respectively. Model 'No-Bot' indicates all three populations maintained constant effective population size, whereas remained models suggest population experienced bottleneck corresponding to population in model name.

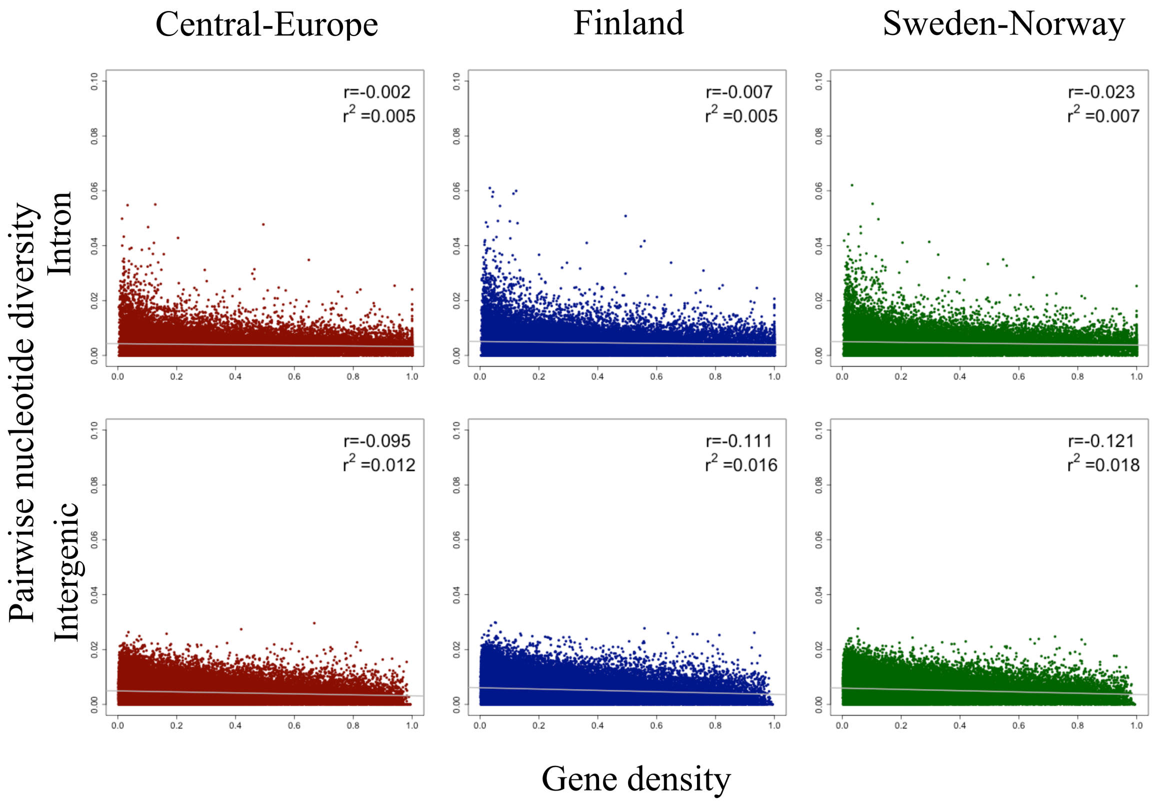


Fig. S4. Scatterplots of intronic nucleotide diversity, intergenic nucleotide diversity versus gene density for Central-Europe population (dark red), Finland population (dark blue), and Sweden-Norway population (dark green) across the *P. abies* genome. The Spearman correlation coefficient (r) and linear regression ($r^2$) are shown in the top right of each plot. Linear regression lines are depicted in grey.

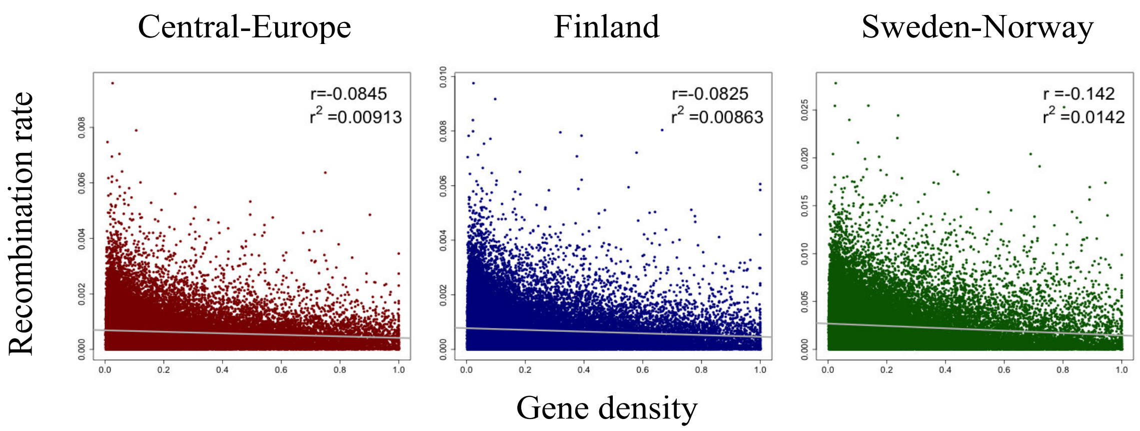


Fig. S5. Scatterplots of the population-scaled recombination rates versus gene density for Central-Europe population (dark red), Finland population (dark blue), and Sweden-Norway population (dark green) across the *P. abies* genome. The Spearman correlation coefficient (r) and linear regression ($r^2$) are shown in the top right of each plot. Linear regression lines are depicted in grey.

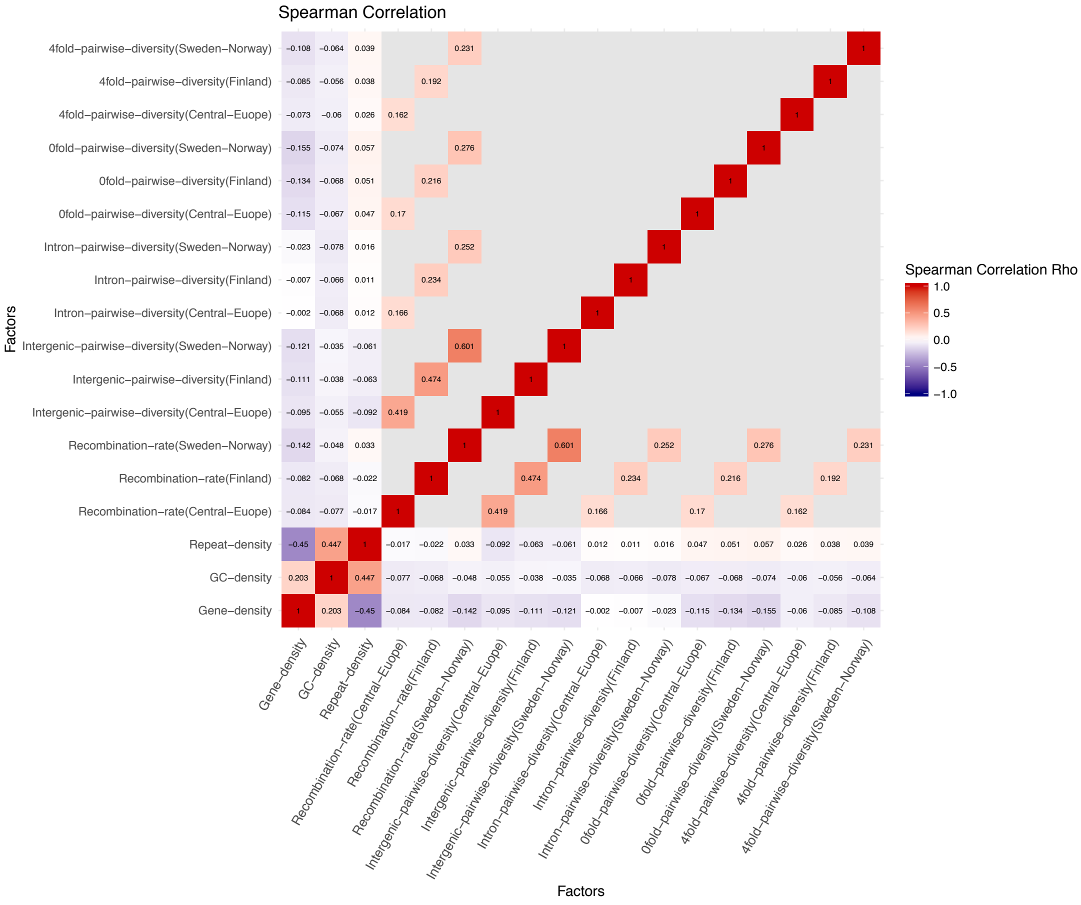


Fig. S6. Heatmap of spearman correlation tests between pairwise factors. Spearman correlation coefficient (r) is shown in each block of pairwise correlation test.
